# Rapid and Quantitative Functional Interrogation of Human Enhancer Variant Activity in Live Mice

**DOI:** 10.1101/2023.12.10.570890

**Authors:** Ethan W. Hollingsworth, Taryn A. Liu, Sandra H. Jacinto, Cindy X. Chen, Joshua A. Alcantara, Evgeny Z. Kvon

## Abstract

Functional analysis of non-coding variants associated with human congenital disorders remains challenging due to the lack of efficient *in vivo* models. Here we introduce dual-enSERT, a robust Cas9-based two-color fluorescent reporter system which enables rapid, quantitative comparison of enhancer allele activities in live mice of any genetic background. We use this new technology to examine and measure the gain- and loss-of-function effects of enhancer variants linked to limb polydactyly, autism, and craniofacial malformation. By combining dual-enSERT with single-cell transcriptomics, we characterize variant enhancer alleles at cellular resolution, thereby implicating candidate molecular pathways in pathogenic enhancer misregulation. We further show that independent, polydactyly-linked enhancer variants lead to ectopic expression in the same cell populations, indicating shared genetic mechanisms underlying non-coding variant pathogenesis. Finally, we streamline dual-enSERT for analysis in F0 animals by placing both reporters on the same transgene separated by a synthetic insulator. Dual-enSERT allows researchers to go from identifying candidate enhancer variants to analysis of comparative enhancer activity in live embryos in under two weeks.

## Introduction

The success of large-scale genome-wide association and whole-genome sequencing studies has shifted the bottleneck of human genetics from identifying sources of genetic variation to mechanistically understanding how such variation contributes to human disease (Claussnitzer et al., 2020; Lappalainen & MacArthur, 2021; Westra & Franke, 2014). Nearly 90% of disease risk-associated variation resides in non-protein coding regions of the human genome (Dong et al., 2023; Edwards et al., 2013; Farh et al., 2015; Turro et al., 2020; F. Zhang & Lupski, 2015). A large fraction of this variation consists of single nucleotide polymorphisms and rare variants that are hypothesized to affect transcriptional enhancers, short non-coding DNA segments that regulate cell-type-specific gene expression (Claringbould & Zaugg, 2021; Gazal et al., 2018; Maurano et al., 2012; Pachano et al., 2022; Ulirsch et al., 2019). Each of these thousands of enhancer variants thus represents a potential entry point for understanding human disease (Claussnitzer et al., 2020; Lappalainen & MacArthur, 2021; Westra & Franke, 2014). However, the physiological effects of the vast majority of these associations remain unknown. Bridging this gap – from non-coding variant to biological mechanism – is currently hindered by a lack of suitable *in vivo* technologies for assessing if and how each human enhancer variant alters gene expression.

A major challenge is that the effects of enhancer variants on gene expression are highly cell-type-specific. For example, a typical gain-of-function enhancer variant can result in ectopic gene expression and cause pathogenic effects in cells where the enhancer is normally inactive (Claussnitzer et al., 2015; Doan et al., 2016; Eufrásio et al., 2020; Kvon et al., 2020; Lewis et al., 2014; Putlyaeva et al., 2017; Turner et al., 2017; Yanchus et al., 2022). Likewise, loss-of-function enhancer variants often result in loss of enhancer activity in one cell type, while in other cell types, its activity is unaffected (Bengani et al., 2021; Bhatia et al., 2015, 2021; Shin et al., 2023; Spieler et al., 2014). These cell-type-specific effects of enhancer variants are difficult to capture with high-throughput methods such as massively-parallel reporter assays (MPRAs) and CRISPR inhibitor/activator screens, both of which are primarily performed *in vitro* (Findlay, 2021; Inoue & Ahituv, 2015; Maricque et al., 2018) or in one tissue (Brown et al., 2022; Capauto et al., 2023; Deng et al., 2023; Lagunas et al., 2023; Patwardhan et al., 2012; White et al., 2013). Transgenic enhancer-reporter assays in mice enable visualization of enhancer activity in the whole animal and are a gold-standard for functionally testing when and where a human enhancer is active *in vivo* (Kothary et al., 1989; Pennacchio et al., 2006; Visel et al., 2006). However, current transgenic mouse reporter assays only allow the assessment of a single enhancer per animal, precluding the direct comparison of multiple enhancer variants. Thus, comparing activities of reference and disease-linked variant alleles requires the generation of a large number of independent transgenic mice to mitigate variation caused by mosaicism and position effects (Kvon et al., 2020; Turner et al., 2017).

Here, we introduce *dual-enSERT* (<u>dual</u>-fluorescent <u>en</u>hancer in<u>SERT</u>ion), a Cas9-based site-specific dual-fluorescent reporter system that enables simultaneous, quantitative visualization of two human enhancer allelic activities in the same transgenic animal, thus overcoming the limitations of standard mouse reporter assays. In *Dual-enSERT-1,* transgenes containing enhancer variants driving *eGFP* or *mCherry* are placed on different alleles of the same safe-harbor location in the mouse genome. In *Dual-enSERT-2*, both enhancer-reporters are placed on the same transgene separated by a synthetic insulator. Using *Dual-enSERT-2,* we were able to visualize and compare two enhancer allelic activities in live F0 mice as soon as eleven days after zygote microinjection. To demonstrate the utility of *dual-enSERT*, we interrogated a panel of new and previously characterized human enhancer variants linked to autism, limb defects, and craniofacial malformation. Beyond quantitative visualization of enhancer activity, *dual-enSERT* captured variant enhancer activity at single-cell resolution. The system thereby allows the nomination of specific molecular pathways as potential mediators of ectopic gene expression caused by disease variants. Our *dual-enSERT* system is thus poised to accelerate enhancer-variant-to-function studies across many congenital disorders.

## Results

### Direct comparison of reference and variant enhancer allele activities *in vivo* with *dual-enSERT-1*

Classical mouse enhancer-reporter assays are based on random integration of the transgene into the genome (Kothary et al., 1989; Kvon, 2015; Pennacchio et al., 2006; Visel et al., 2006; Zakany et al., 1988). Although conventional mouse transgenesis is the current gold standard for visualization of enhancer activity *in vivo*, it suffers from variation due to position effects and therefore requires generation of a large number of transgenic animals to reproducibly assess enhancer activity (Fakhouri et al., 2014; Lettice et al., 2008; Turner et al., 2017). Enhancer-reporter assays based on the integration of a transgene into a safe-harbor location of the genome overcomes the problem of position effects (Kvon et al., 2020; Tasic et al., 2011). However, due to mosaicism and inter-embryo variability, these site-specific assays also require analysis of a large number of transgenic animals, especially for detecting the subtle effects of enhancer mutations (Kvon et al., 2020; Osterwalder et al., 2022; Snetkova et al., 2021). To overcome these limitations, we developed *dual-enSERT-1*, a transgenic approach based on highly-efficient Cas9-mediated integration of enhancers driving fluorescent reporters into the H11 safe-harbor integration site (Hippenmeyer et al., 2010). One enhancer allele is placed upstream of an *eGFP* reporter and the second enhancer allele is placed upstream of an *mCherry* reporter followed by Cas9-mediated integration of each transgene into the H11 locus (**Fig. 1A**). With *dual-enSERT-1* we achieved an average transgenic targeting efficiency of 57% across all single-reporter constructs tested in this study, and all insertions were germline transmissible (**Fig. 1B**).

**Figure 1:**
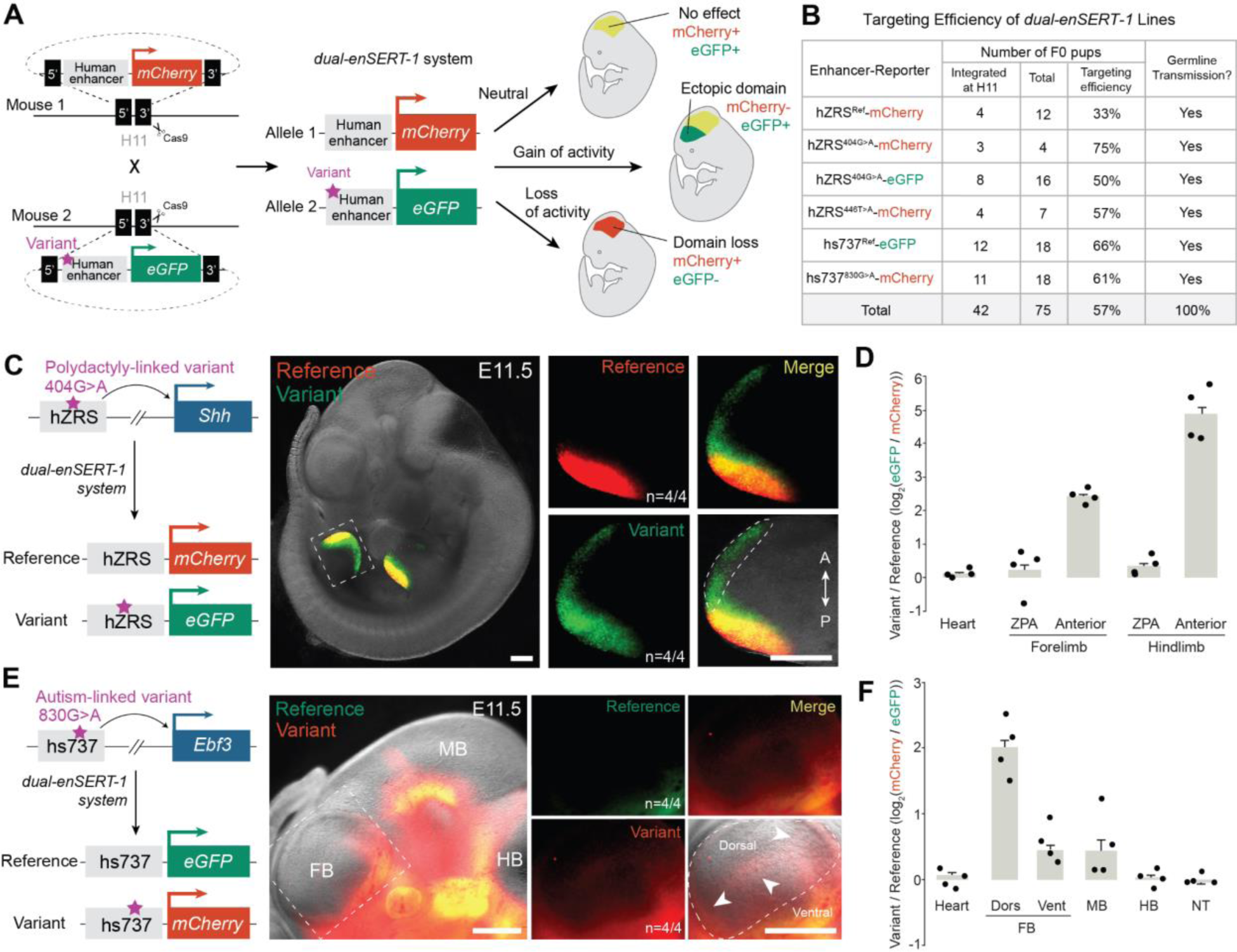
Simultaneous comparison of human reference and variant enhancer activities. (**A**) Schematic overview of the *dual-enSERT-1* strategy and potential readouts for different types of disease-linked enhancer variants. (**B**) Transgenic targeting efficiencies of all *dual-enSERT-1* constructs generated in this study. (**C**) Representative images of transgenic hZRS^ref^-*mCherry*/hZRS^404G>A^-*eGFP* embryos at E11.5. A close-up of the hindlimb with separate and merged channels and outline depicting gain of enhancer activity (green channel) in the anterior domain are shown. A, anterior; P, posterior. (**D**) Plots quantifying fold-change (log2) difference in reporter intensity between variant and reference ZRS alleles. Two-sided paired t-tests: Forelimb ZPA, *P =* ns; Forelimb Anterior, *P =* 0.00012; Hindlimb ZPA, *P =* ns; Hindlimb Anterior, *P =* 2.40E-05. (**E**) Representative images of transgenic hs737^ref^-*eGFP*/hs737^830G>A^-*mCherry* embryos at E11.5. A close-up of the forebrain with separate and merged channels are shown. White outline and arrows highlight ectopic areas with gain in enhancer activity (red channel). (**F**) Plots quantifying fold-change (log2) difference in reporter intensity between variant and reference hs737 alleles. Two-sided paired t-tests: Ventral Forebrain (FB), *P* = ns; Dorsal Forebrain, *P* = 0.0020; Midbrain (MB), *P* = ns; Hindbrain (HB), *P* = ns; Neural Tube (NT), *P* = ns. Data represented as mean ± SEM for all plots. All scale bars, 500 μm.

We first assessed whether we could detect and quantify the effects of non-coding variants on enhancer activity with *dual-enSERT-1* by testing a previously characterized pathogenic allele of the ZRS (zone of polarizing activity (<u>Z</u>PA) <u>R</u>egulatory <u>S</u>equence, also known as MFCS1) enhancer of *Sonic hedgehog (Shh)*. Single nucleotide variants in the ZRS cause congenital limb malformations, most typically preaxial polydactyly, in humans, cats, chicken, and mice (Heutink et al., 1994; Kvon et al., 2020; Lettice et al., 2003). ZRS variants implicated in preaxial polydactyly cause ectopic *Shh* expression in the anterior portion of the developing limb bud, leading to erroneous digit outgrowth similar to human patients (Kvon et al., 2020; Lettice et al., 2003). We created two stable transgenic mouse lines, one with the human reference ZRS allele driving *mCherry* (hZRS^ref^-*mCherry)* and a second line with a previously characterized pathogenic ZRS allele containing the polydactyly-linked 404G>A variant driving *eGFP* (hZRS^404G>A^-*eGFP*) (**Fig. 1C**). To visualize reference and variant ZRS enhancer activities simultaneously, we crossed these mouse lines to generate two-color *dual-enSERT-1* embryos. In these hZRS^ref^-*mCherry*/hZRS^404G>A^-*eGFP* transgenic embryos, mCherry fluorescence was only detected in the zone of polarizing activity (ZPA) of fore- and hindlimb buds, matching the location of normal *Shh* expression (Riddle et al., 1993). EGFP expression driven by the ZRS^404G>A^ variant allele was detected in the ZPA at comparable levels with mCherry expression at embryonic day 11.5 (Forelimb, *P* = ns; Hindlimb, *P* = ns). EGFP expression driven by the ZRS^404G>A^ variant allele also extended into the ectopic anterior domain of the limb bud in all examined embryos, mimicking *Shh* misexpression in mouse embryos containing pathogenic mutations in the ZRS (5.4-fold difference in Forelimb, *P* = 0.00012; 32-fold difference in Hindlimb, *P* = 2.40E-05; **Fig. 1C, D**). These results indicate that *dual-enSERT-1* can robustly detect and quantify changes in limb enhancer activity caused by pathogenic ZRS variants.

We next asked whether *dual-enSERT-1* could be used to study human non-coding variants linked to congenital disorders affecting other tissues. We focused on the hs737 enhancer of *Ebf3*, a region where several independent rare variants have been identified in patients with autism spectrum disorder and intellectual disability (Padhi et al., 2021; Turner et al., 2017). We generated one mouse line in which the human reference hs737 allele drives *eGFP* (hs737^ref^-e*GFP*) and a second line in which an 830G>A variant allele, identified in a patient with autism, drives *mCherry* (hs737^830G>A^-*mCherry*) (**Fig. 1E**). We then bred these mouse lines and examined reporter gene expression in E11.5 embryos. Live imaging revealed comparable levels of eGFP and mCherry fluorescence in the ventral forebrain, midbrain, hindbrain, and neural tube (Ventral Forebrain, *P* = ns; Midbrain, *P* = ns; Hindbrain, *P* = ns; Neural Tube, *P* = ns; **Fig. 1E, F**). mCherry expression driven by the hs737^830G>A^ variant allele also extended to the dorsal part of forebrain in all examined hs737^ref^-*eGFP*/hs737^830G>A^-*mCherry* embryos (4.2-fold difference in Dorsal Forebrain, *P* = 0.0020; **Fig. 1E, F**). These results are consistent with previous observations of ectopic forebrain activity using non-quantitative LacZ-based transgenic assays (Turner et al., 2017). Taken together, these results indicate that *dual-enSERT-1* can robustly and reproducibly visualize activities of two enhancer alleles and detect the quantitative effects of non-coding variants on enhancer activity in multiple tissues in live mice.

### Comparative functional assessment of independent enhancer variants

We next tested the ability of *dual-enSERT-1* to simultaneously visualize and compare the effects of independent enhancer variants in live mice, including previously uncharacterised variants. Human genetics studies often identify disease-linked hotspots in which multiple rare variants affect the same enhancer (Chatterjee et al., 2016; Dunning et al., 2016; Kvon et al., 2020; Pomerantz et al., 2009; Ulirsch et al., 2019; X. Zhang et al., 2012). Due to the low reproducibility of canonical enhancer-reporter assays (Kvon et al., 2020), it is challenging to determine whether these independent disease-linked variants cause loss or gain of activity in the same cell types. This knowledge is critical to determine the pathogenic consequences of existing and newly identified genetic variants that map to a previously characterised non-coding locus. For example, 22 different rare human point mutations in the ZRS enhancer have been identified in patients with polydactyly (Kvon et al., 2020; Lettice et al., 2003). Despite extensive work on this enhancer, it is unknown if these independent mutations result in ectopic gene expression in the same or different cell populations of the limb bud. To address this question, we generated a transgenic mouse line in which a new ZRS^446T>A^ variant allele identified in a family with preaxial polydactyly drives *mCherry* (hZRS^446T>A^-*mCherry*) (Xu et al., 2020). The 446T>A variant is hypothesized to create a *de novo* activator binding site, but its effect on *in vivo* ZRS enhancer activity is unknown^55^. In contrast, the well-characterised 404G>A variant disrupts a repressor binding site and causes ectopic reporter expression in the anterior limb bud mesenchyme (Lettice et al., 2017, 2003). To visualize hZRS^404G>A^ and ZRS^446T>A^ variant allele activities simultaneously, we bred these mouse lines to generate two-color transgenic embryos. In these hZRS^446T>A^-*mCherry*/hZRS^404G>A^-e*GFP* transgenic embryos, mCherry and eGFP were detected in a highly overlapping pattern in the ZPA and anterior limb bud mesenchyme at E11.5 (ZPA, *P* = ns; Anterior, *P* = ns; **Fig. 2A and S1**). The overlap in ectopic activity in the anterior limb bud mesenchyme was visually and quantitatively indistinguishable from the overlap observed in hZRS^404G>A^-*mCherry*/hZRS^404G>A^-e*GFP* transgenic embryos in which *eGFP* and *mCherry* were driven by the same ZRS^404G>A^ variant allele (**Fig. 2B and S1**).

**Figure 2:**
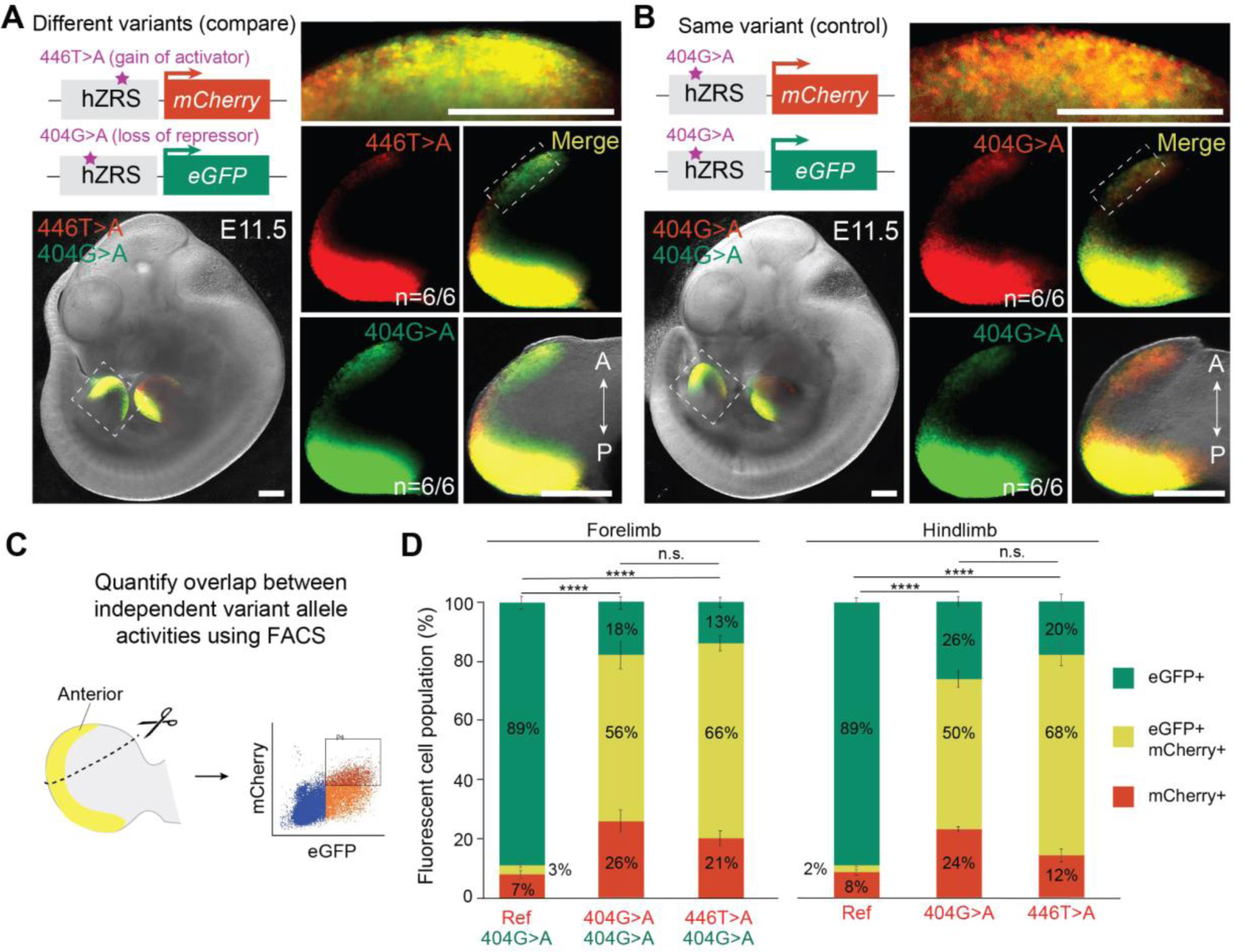
Comparison of the effects of independent human-disease-linked variants in the ZRS enhancer. **(A)** Representative image of hZRS^446T>A^-*mCherry*/hZRS^404G>A^-*eGFP* embryo at E11.5. Panels on right show high-resolution images of hindlimb bud and its anterior domain (above), as marked by outlined boxes. **(B)** Sample image of E11.5 hZRS^404G>A^-*mCherry*/hZRS^404G>A^-*eGFP* embryo. Panels on right show high-resolution images of hindlimb bud and anterior domain (above), as marked by outlined boxes. **(C)** FACS-based quantification of the overlap between *mCherry*- and *eGFP*-expressing cells in the anterior domain of limb buds. (**D**) Plots depicting population distribution of eGFP+, eGFP+ mCherry+, and mCherry+ cells in anterior domain of fore- and hindlimbs. Genotypes of *dual-enSERT-1* embryos are labeled along the x-axis, colored according to the downstream reporter gene. Fisher exact test; *P* < 0.0001 (****). Data represented as mean ± SEM for plots. All scale bars, 500 μm.

To quantify the overlap between ectopic activities of different ZRS alleles at cellular resolution, we used fluorescent-associated cell sorting (FACS) to isolate double-positive cells from anterior limb bud mesenchyme. We dissected the anterior portion of fore- and hindlimb buds from E11.5 *dual-enSERT-1* embryos followed by cell dissociation and FACS (**Fig. 2C and S1**). As a negative control, we used transgenic mice in which the ZRS^404G>A^ variant allele was driving *eGFP* and the ZRS reference allele was driving *mCherry* (hZRS^ref^-*mCherry*/hZRS^404G>A^-*eGFP*) with the expectation that only eGFP+ cells should be present in the anterior limb bud cell population (**Fig. 1B**). Indeed, 89% of fluorescent cells sorted from anterior limb buds of hZRS^ref^-*mCherry*/hZRS^404G>A^-*eGFP* embryos were eGFP+/mCherry- and only 3% were eGFP+/mCherry+, confirming allele-specific ectopic expression of a variant ZRS^404G>A^ allele (**Fig. 2D**). We next examined hZRS^404G>A^-*mCherry*/hZRS^404G>A^-*eGFP* transgenic embryos carrying the ZRS^404G>A^ variant allele driving both colors to account for stochasticity in reporter expression and any differences between fluorophores (**Fig. 2A**). 50% (in forelimbs) to 56% (in hindlimbs) of fluorescent cells in anterior limb buds were eGFP+/mCherry+, indicating a significant overlap between eGFP and mCherry ectopic expression driven by the same 404G>A variant (**Fig. 2D**). Finally, we examined hZRS^446T>A^-*mCherry*/hZRS^404G>A^-*eGFP* transgenic embryos in which *eGFP* and *mCherry* were driven by different ZRS variants (**Fig. 2B**). 66% (in forelimbs) to 68% (in hindlimbs) of fluorescent cells in anterior limb buds were eGFP+/mCherry+ (**Fig. 2D**). This fraction of double positive anterior limb bud cells was not significantly different from the fraction of double-positive cells in hZRS^404G>A^-*mCherry*/hZRS^404G>A^-*eGFP* transgenic embryos. These results show that independent 404G>A and 446T>A variants cause highly overlapping ectopic expression, indicating that a shared genetic mechanism may underlie ZRS enhancer misregulation. Altogether, these results demonstrate the power of *dual-enSERT-1* for direct comparison of independent enhancer variants.

### Pathogenic enhancer variant activity at single-cell resolution

We next asked whether ectopic activity caused by enhancer variants can be quantitatively assigned to specific cell types *in vivo* using *dual-enSERT-1*. Such information coupled with gene expression profiling can potentially reveal which genetic pathways lead to ectopic gene expression upon enhancer misregulation. We focused on the pathogenic hZRS^404G>A^ variant allele for which the mechanism of ectopic *Shh* expression in the limb bud is not known^41^ (**Fig. 1C**). We dissected a hindlimb bud from an E11.5 hZRS^ref^-*mCherry*/hZRS^404G>A^-*eGFP* transgenic embryo and performed 10X single-cell RNA-sequencing (scRNA-seq) (**Fig. 3A and S2**). Because scRNA-seq often suffers from gene dropout (Kharchenko et al., 2014; Qiu, 2020), we adopted a nested PCR strategy to amplify *mCherry* and *eGFP* transcripts in our barcoded libraries (**Methods; Fig. 3A**) (Pollina et al., 2023). After quality control, we obtained scRNA-seq profiles for over 15,000 cells, which clustered into thirteen distinct cell types, including a large mesenchymal cluster defined by the specific expression of well-known marker genes (**Fig. 3B and S2**) (Desanlis et al., 2020; Yokoyama et al., 2017). For the hZRS^ref^ allele, *mCherry* expression was almost exclusively restricted to the distal posterior mesenchyme, matching expression of *Shh* (**Fig. 3C-E**). By contrast, *eGFP* expression driven by the hZRS^404G>A^ allele was also detected in distal middle, anterior, and proximal anterior mesenchymal cells, matching the distribution of EGFP fluorescence in live embryos (**Fig. 3C, D**).

**Figure 3:**
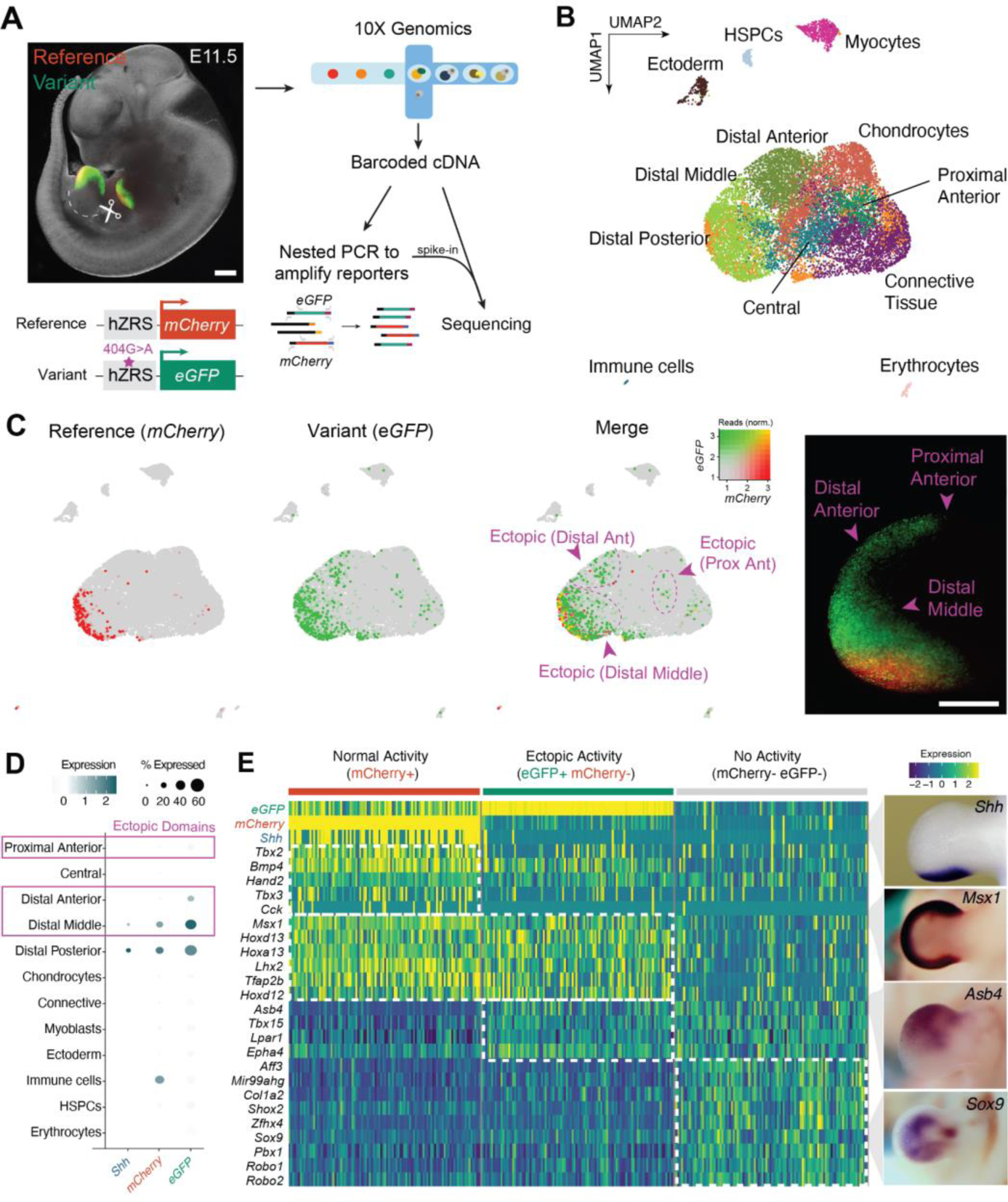
Molecular characterization of pathogenic enhancer variant activity at single-cell resolution. (**A**) Single-cell transcriptomic profiling of a hindlimb from transgenic hZRS^ref^-*mCherry*/hZRS^404G>A^-*eGFP* mice at E11.5. A nested PCR strategy was used to amplify *mCherry* and *eGFP* reporter transcripts (see **Methods** for details). (**B**) UMAP plot showing cell clusters that are found in the E11.5 mouse hindlimb bud. Mesenchymal cell clusters defined by expression of spatially-defined genes are shown. (**C**) Feature plots showing *mCherry* and *eGFP* expression with overlapping cells marked in yellow. Areas of ectopic eGFP expression are highlighted in magenta. (**D**) A dot plot depicting percentage and expression of cells expressing e*GFP*, *mCherry*, and *Shh* within each cell cluster. Cell clusters with ectopic *eGFP* expression are highlighted. (**E**) Heatmap of differentially expressed genes between mCherry+ (normal ZRS activity), eGFP+ mCherry-(ectopic ZRS activity), and mCherry-eGFP-(ZRS is inactive) cell subpopulations. Spatial distribution of representative marker genes in E11.5 limb buds are shown on the right. Images have been reproduced with permission from Embrys database (http://embrys.jp). Scale bars, 500 μm.

To characterize cell populations in which reference and variant enhancer alleles are active, we examined mCherry+/eGFP-(‘normal’ ZRS activity), eGFP+/mCherry-(‘ectopic’ ZRS activity) and mCherry-/eGFP-(inactive) subpopulations. We then performed unbiased differential gene expression analysis between these cell populations to identify candidate genetic pathways linked to normal and ectopic *Shh* expression. Among the cells with ectopic ZRS activity, the most-enriched genes were a number of anterior-specific genes like *Asb4* and *Epha4.* This pattern was distinct from cells with normal enhancer activity which were enriched for *Shh* and genes critical for establishing *Shh* expression in the ZPA, such as *Hand2* (**Fig. 3E**) (Osterwalder et al., 2014). By contrast, inactive cells expressed chondrocyte-specifying transcription factors like *Shox2* and *Sox9* (**Fig. 3E**) (Akiyama et al., 2002; Yu et al., 2007). Notably, both normal and ectopic ZRS domains showed a surprising similarity in their transcriptional profiles with enrichment of several distal mesenchymal transcription factors, including *Msx1*, *Hoxd12*, *Hoxd13*, *Hoxa13*, and *Lhx2* (**Fig. 3E**). This transcriptomic overlap among subpopulations may explain the propensity for mutations in the ZRS to cause ectopic *Shh* expression in anterior cells. Taken together, these results implicate candidate pathways in ectopic ZRS activity and highlight how the capture of variant allele-labeled cells via *dual-enSERT-1* opens opportunities for mechanistic study of pathogenic enhancer misregulation.

### ZRS enhancer bypasses three copies of the chicken *β*-globin insulator

A limitation of the *dual-enSERT-1* system is the ∼6 month time required to obtain two-color F2 embryos. This protracted timeframe limits the number of enhancer variants that can be rapidly tested in mice. To overcome this bottleneck, we constructed single transgenes containing *mCherry* and *eGFP* reporters driven by different enhancer alleles in divergent orientations. With this bicistronic system, henceforth referred to as *dual-enSERT-2.0*, an injection of a single construct would yield two-color F0 embryos in as little as ten days (**Fig. 4A**). As a proof of principle, we placed the hZRS^ref^ allele upstream of *mCherry* and hZRS^404G>A^ variant allele upstream of *eGFP* all within the same construct. To prevent any potential cross-activation between enhancer alleles, we separated them using three copies of a previously characterised chicken *β*-globin insulator, 5’-HS4. 5’-HS4 is widely used for its robust ability to block enhancer-promoter activation in the genome (Bungert et al., 1995; Huang et al., 2021; Yusufzai & Felsenfeld, 2004) and in the context of a zebrafish transgene (Bhatia et al., 2021). We also placed an additional three copies of the 5’-HS4 insulator (3xHS4) into the plasmid backbone to prevent cross-activation between different copies of the transgene in the event of multi-copy integrations at the H11 landing site (Kvon et al., 2020) (**Fig. 4B and S3**). We injected the resulting hZRS^ref^-*mCherry*/3xHS4/hZRS^404G>A^-*eGFP* bicistronic construct into mouse zygotes and collected transgenic embryos at E11.5. Surprisingly, we detected robust mCherry and eGFP expression in anterior cells in all examined transgenic embryos, indicating that variant ZRS alleles can simultaneously activate *eGFP* and *mCherry* reporter genes in this transgene context (5/5 of embryos with a single-copy transgene integration at H11, 11/11 of embryos with multi-copy transgene integration at H11, **Fig. 4B and Table S1**). Quantitatively, we found no difference in fluorescent intensity across the anterior and ZPA regions of the fore- and hindlimb (Forelimb ZPA, *P* = ns; Forelimb Anterior, *P* = ns; Hindlimb ZPA, *P* = ns; Hindlimb Anterior, *P* = ns; **Fig. 4C**). These results show that three copies of the 5’-HS4 insulator are insufficient to insulate ZRS alleles from cross-activating reporter genes (**Fig. 4D**).

**Figure 4:**
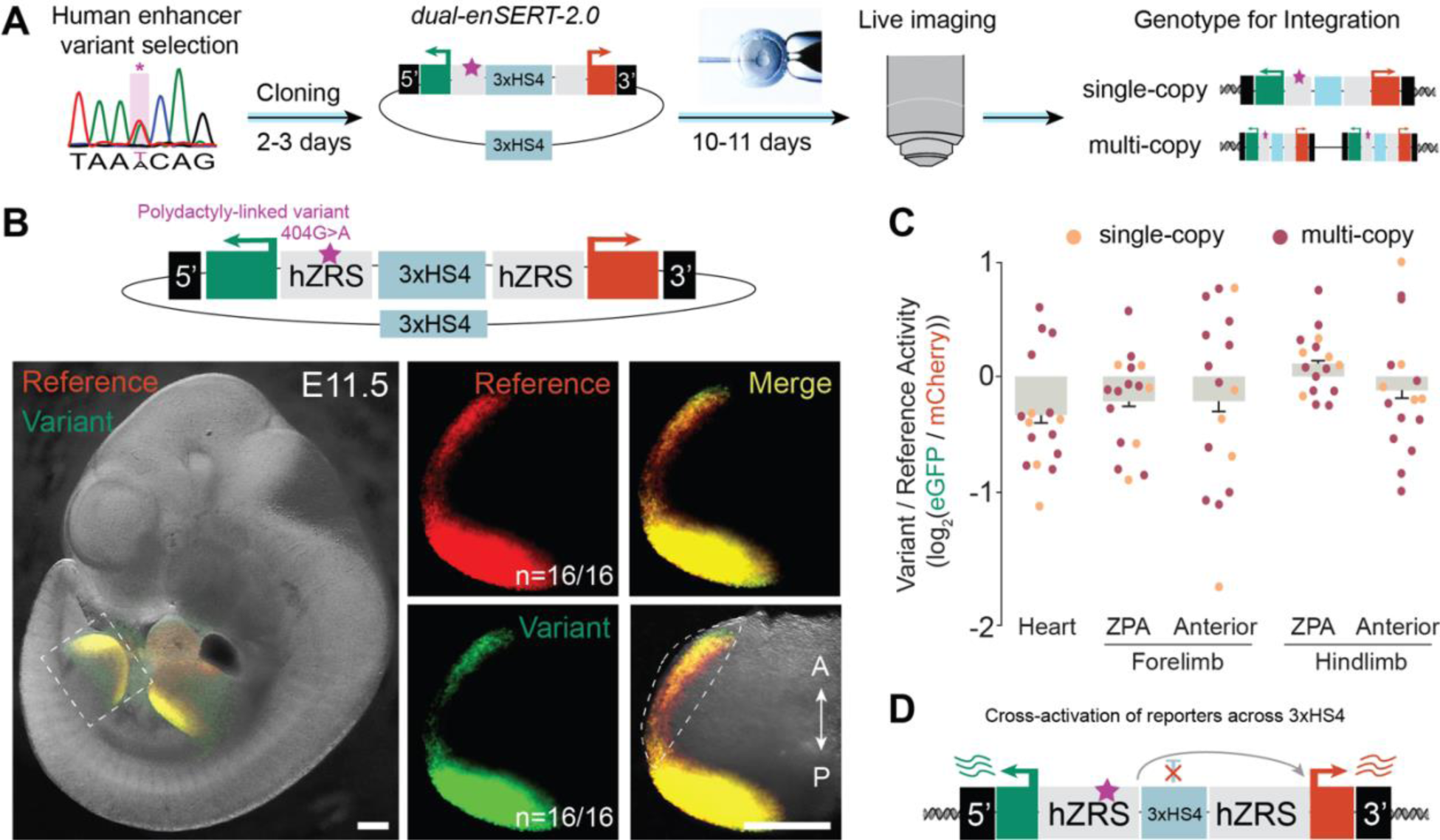
Three copies of the chicken β-globin insulator fail to prevent *mCherry* cross-activation by the variant ZRS allele. (**A**) Schematic design of the *dual-enSERT-2.0* system with enhancer alleles driving *eGFP* or *mCherry* are placed on the same transgene and separated by three copies of the chicken β-globin insulator. (**B**) Sample image of hZRS^ref^-*mCherry*/3xHS4/hZRS^404G>A^-*eGFP* embryo at E11.5. Panels on right show an expanded view of hindlimb. Scale bars, 500 μm. (**C**) Fold-change (log2) quantification variant to reference allele fluorescent reporter intensity in mouse embryos injected with hZRS^ref^-*mCherry*/3xHS4/hZRS^404G>A^-*eGFP* plasmid. Data points representing single-(n=5) and multi-copy (n=11) integrants are in orange and maroon, respectively. Two-sided paired t-tests: Forelimb ZPA, *P =* ns; Forelimb Anterior, *P =* ns; Hindlimb ZPA, *P =* ns; Hindlimb Anterior, *P =* ns. (**D**) Schematic summary depicting lack of insulation by three copies of the 5’-HS4 between enhancer-reporter transgenes.

### A strong synthetic insulator prevents reporter cross-activation in a single-copy but not multi-copy transgene

We next tested if a stronger insulator could prevent enhancer-reporter cross-activation in our F0 *dual-enSERT-2.0* transgene. Previous studies suggest that a number of functional elements, such as CTCF binding sites within the insulator, positively correlate with its enhancer-blocking activity (Bell et al., 1999; Huang et al., 2021; Jia et al., 2020; Yusufzai et al., 2004). There is also evidence that insulator function is transcription factor-specific, i.e. some insulators work with one type of enhancer but not with another (Hong et al., 2022; Ribeiro-Dos-Santos et al., 2022). We therefore created a strong synthetic insulator (SI) by fusing multiple copies of three of the most well-studied vertebrate insulators: A2 (two copies), ALOXE3, and 5’-HS4 (two copies) (Bungert et al., 1995; Liu et al., 2015; Raab et al., 2012) (**Fig. S3**). Importantly, these three insulators utilize different mechanisms for insulation: A2 and 5’-HS4 depend on CTCF while ALOXE3 relies on two B-boxes that recruit RNA polymerase III (Farrell et al., 2002; Liu et al., 2015; Schramm & Hernandez, 2002). In principle, the combination of these three insulators should block most enhancers from activating promoters. We placed one copy of this synthetic insulator between hZRS^ref^ and 404G>A variant alleles and one copy in the plasmid backbone of this new *dual-enSERT-2.1* vector (**Fig S3**). We injected ZRS^ref^-*mCherry*/SI/ZRS^404G>A^-*eGFP* plasmid into mouse zygotes together with Cas9 RNPs targeting the H11 locus, and imaged F0 embryos at E11.5. In embryos with a single-copy transgene integration at H11 locus, ZRS^ref^-driven mCherry fluorescence was restricted to the ZPA while ZRS^404G>A^-driven eGFP also extended to the anterior margin of the limb bud in all examined embryos, mimicking the results of *dual-enSERT-1* in which enhancer-reporters were placed on separate H11 alleles (3/3 embryos; Forelimb ZPA, *P* = ns; 1.9-fold difference in Forelimb Anterior, *P* = 0.0229; Hindlimb ZPA, *P* = ns; 2.1-fold difference in Hindlimb Anterior, *P* = 0.00898; **Fig. S3** and **Fig. 1B**). These results indicate that in the *dual-enSERT-2.1* system, the synthetic insulator can successfully block ZRS-reporter cross-activation at the H11 locus. Interestingly, in embryos with multiple copies of a transgene separated by a synthetic insulator at the H11 locus, both anterior and posterior limb buds showed robust mCherry and EGFP expression in all examined embryos (8/8 embryos; Forelimb ZPA, *P* = ns; Forelimb Anterior, *P* = ns; Hindlimb ZPA, *P* = ns; Hindlimb Anterior, *P* = ns; **Fig. S3**). These results indicate that the ZRS can bypass the synthetic insulator in the context of a tandem transgene with multiple enhancer-reporters flanked by synthetic insulators (**Fig. S3**). Therefore, a synthetic insulator-based *dual-enSERT-2.1* setup can only discriminate between enhancer allele activities if a single copy of the bicistronic transgene is integrated at the H11 landing site (**Fig. S5**).

### *Dual-enSERT-2.2* allows rapid visualization of two enhancer allele activities using a single-copy transgene

To optimize the efficiency of *dual-enSERT-2.1*, we developed *dual-enSERT-2.2* by maximizing the number of single-copy integrants at the H11 landing site (**Fig. 5A**). Recent work in zebrafish and mice has shown that addition of biotinylated nucleotides to the ends of donor DNA prevents concatemer formation during Cas9-mediated homology-directed repair (Gutierrez-Triana et al., 2018; Medert et al., 2023). To test if addition of biotin (B) results in preferential single-copy transgene integration at the H11 locus, we added biotinylated nucleotides to the ends of a linearised F0 dual-enSERT vector carrying the human reference and the 404G>A ZRS variant alleles (**Methods**). We injected this B-hZRS^ref^-*mCherry*/SI/hZRS^404G>A^-*eGFP*-B construct into mouse zygotes together with Cas9 RNPs and imaged the F0 embryos eleven days later. mCherry fluorescence was restricted to the ZPA, while the ZRS 404G>A variant-driven eGFP expression extended into the anterior limb bud in all embryos with transgene integration at the H11 locus (5/5 embryos, Forelimb ZPA, *P* = ns; 1.3-fold difference in Forelimb Anterior, *P* = 0.0349; Hindlimb ZPA, *P* = ns; 1.5-fold difference in Hindlimb Anterior, *P* = 5.20E-05; **Fig. 5B, C and Table S1**). Moreover, genotyping confirmed that most transgenic embryos (n=5/6 embryos) contained a single-copy transgene at the H11 locus and lacked concatemers (**Fig. 5B, C and Table S1**). This result indicates that addition of biotinylated nucleotides results in preferential single-copy integration of a bicistronic transgene and enables discrimination between enhancer allele activities in a straightforward manner.

**Figure 5:**
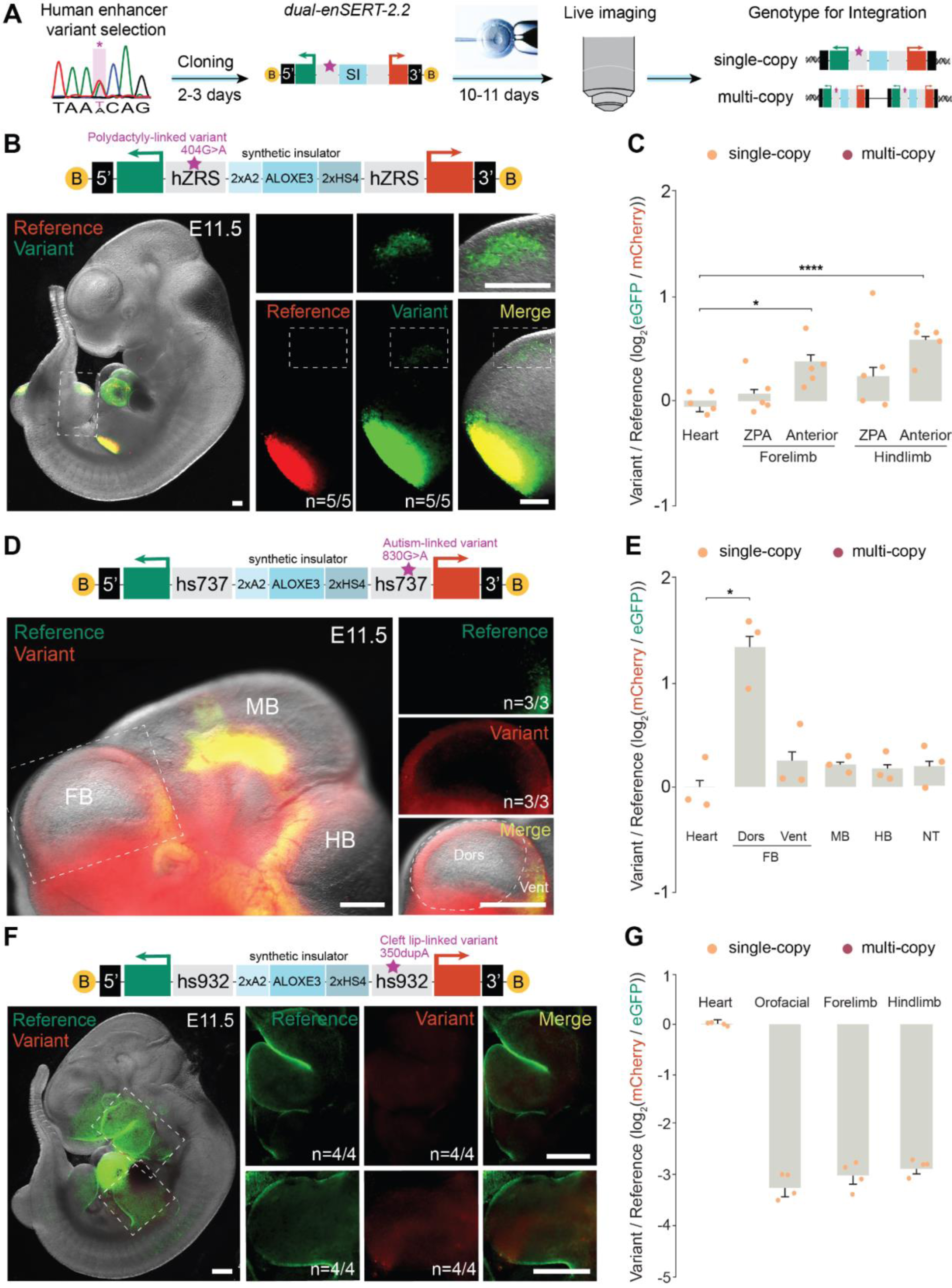
An F0-based *dual-enSERT-2.2* system for rapid testing of human enhancer variant activity. (**A**) Schematic design of the *dual-enSERT-2.2* system with biotinylated nucleotides attached to the end of donor fragments for preferential single-copy integration and a synthetic insulator (SI) to prevent cross activation between transgenes. (**B**) Fluorescent image of B-hZRS^ref^-*mCherry*/SI/hZRS^404G>A^-*eGFP*-B whole embryo at E11.5 with close-up images of hindlimb and anterior domain on right as marked by white dotted boxes. B, biotin. Scale bars, 250 μm. (**C**) Plots quantifying fold-change (log2) difference in reporter intensity between variant and reference alleles by tissue in B-hZRS^ref^-*mCherry*/SI/hZRS^404G>A^-*eGFP*-B embryos. Data points representing single-copy (n=5) integrants are in orange. Two-sided paired t-tests: Forelimb ZPA, *P =* ns; Forelimb Anterior, *P =* 0.0349; Hindlimb ZPA, *P =* ns; Hindlimb Anterior, *P =* 5.20E-05. (**D**) Sample fluorescent image of a B-hs737^ref^-*eGFP*/SI/hs737^830G>A^-*mCherry*-B embryo at E11.5 with high-resolution image of forebrain on right. White outline highlights ectopic areas with the gain in enhancer activity (red channel). B, biotin. Scale bars, 500 μm. (**E**) Plot depicting quantification of fold-change (log2) difference in variant-reference reporter intensity by tissue in B-hs737^ref^-*eGFP*/SI/hs737^830G>A^-*mCherry*-B embryos. Two-sided paired t-tests: Dorsal (Dors) Forebrain, *P* = 0.044; Ventral (Vent) Forebrain (FB), *P* = ns; Midbrain (MB), *P* = ns; Hindbrain (HB), *P* = ns; Neural Tube (NT), *P* = ns. Data represented as mean ± SEM for all plots. (**F**) Representative image of B-hs932^350dupA^-*mCherry*/SI/hs932^ref^-*eGFP*-B embryo, with expanded views on right to highlight orofacial region and limbs at E11.5, as marked by white dotted boxes. B, biotin. Scale bars, 500 μm. (**G**) Quantitative plot for fold-change (log2) difference in variant-reference fluorescent reporter intensity for heart, orofacial region, and limbs from B-hs932^ref^-*eGFP*/SI/hs932^350dupA^-*mCherry*-B embryos. Data points representing single-copy integrants (n = 4) are in orange. Two-sided paired t-tests: Orofacial, *P =* 0.00023; Forelimb, *P* = 0.00047; Hindlimb, *P =* 0.00035. Data represented as mean ± SEM for all plots.

To test if *dual-enSERT-2.2* could rapidly detect the quantitative effects of other disease-linked non-coding variants, we returned to the autism- and intellectual disability-linked hs737/*Ebf3* locus (**Fig. 1E, F**). We placed the hs737^ref^ allele upstream of *eGFP* and the hs737^830G>A^ variant allele upstream of *mCherry*. We then linearised, biotinylated, and injected the resulting B-hs737^ref^-*eGFP*/SI/hs737^830G>A^-*mCherry*-B construct into mouse zygotes with Cas9 RNPs. Eleven days later, live imaging revealed increased hs737^830G>A^-driven mCherry in the dorsal forebrain (2.6-fold, *P* = 0.043) while the ventral forebrain (*P* = ns), midbrain (*P* = ns), hindbrain (*P* = ns), and neural tube (*P* = ns) showed no difference between EGFP and mCherry reporter intensity (**Fig. 5D, E**, and **Table S1**). These results reproduce our earlier results using *dual-enSERT-1*, but in much shorter time frame (11 days vs. 6 months).

Finally, we asked if *dual-enSERT-2.2* could be employed to study human disease variants that cause loss of enhancer activity. We focused on the previously characterised rare non-coding 350dupA mutation at the *IRF6* locus that is linked to cleft lip formation (Fakhouri et al., 2014). 350dupA is a single A duplication at position 350 of the hs932 face enhancer of *IRF6* (also known as MCS9.7). We created a bicistronic vector with the hs932 reference allele driving *eGFP* and the hs932^350dupA^ variant allele driving *mCherry* separated by the synthetic insulator (**Fig. 5F**). We biotinylated the ends of the linearised vector and injected the resulting B-hs932^ref^-*eGFP*/SI/hs932^350dupA^-*mCherry*-B construct together with Cas9 RNPs into zygotes. Imaging of transgenic mouse embryos at E11.5 revealed strong eGFP expression driven by the reference hs932 allele in the orofacial and limb ectoderm, as previously reported using LacZ-based transgenesis (Fakhouri et al., 2014) (4/4 embryos, **Fig. 5F** and **Table S1**). By contrast, we found no mCherry fluorescence driven by the variant hs932^350dupA^ allele in all examined transgenic embryos, indicating a near complete loss of enhancer activity (10-fold in orofacial, *P* = 0.00023; 8.5-fold in forelimb, *P* = 0.00047; 7.7-fold in hindlimb, *P* = 0.00035; **Fig. 5F, G**). Crucially, 93% of all enhancer-reporters integrated at the H11 locus across the three tested lines were single-copy transgenes allowing easy interpretation of imaging results (**Table S1**). Altogether, these results demonstrate the utility of *dual-enSERT-2.2* for rapidly detecting and quantifying gain- and loss-of-activity for disease-linked enhancer variants *in vivo*.

## Conclusions

Realising the functional and therapeutic potential promised by large-scale human genomics studies depends on our ability to test the *in vivo* effects of candidate non-coding variants linked to human disease. However, no method currently exists for direct comparative testing of enhancer variant activity in a live mammal. In this study, we directly addressed this unmet need by developing *dual-enSERT*, a rapid, quantitative, and cost-effective method for simultaneous comparison of human enhancer allele activities in live mice. By applying *dual-enSERT* to a diverse panel of previously characterised and new disease-linked enhancer variants, we demonstrated its widespread compatibility across diverse congenital disorders and organ systems. *Dual-enSERT* can also be easily combined with single-cell technologies, as demonstrated by our data suggesting that independent non-coding mutations cause polydactyly through potentially shared genetic mechanisms.

While developing *dual-enSERT-2*, we found that three copies of the widely used 5′-HS4 chicken β-globin insulator unexpectedly failed to block communication between the ZRS enhancer and the *hsp68* minimal promoter (**Fig. 4**). These results support previous observations that insulator function can be locus- or enhancer-specific (Hong et al., 2022; Ribeiro-Dos-Santos et al., 2022; Uchida et al., 2011) and suggest that caution should be taken when using individual insulators to protect transgenes in genome engineering applications. Our synthetic insulator, consisting of three different previously characterised insulators, can effectively block ZRS, hs737, and hs932 enhancer-promoter communication in the context of a transgene and has the potential to work in a wider number of genomic contexts.

Beyond studying disease-linked enhancer variants, *dual-enSERT* can be used for evolutionary developmental biology studies. Sequence divergence in enhancers is hypothesized to be a major driver of morphological and functional evolution (Long et al., 2016; Rebeiz & Tsiantis, 2017); however pinpointing and functionally testing the causal regulatory regions has been challenging. With *dual-enSERT*, the activity of candidate evolution-driving enhancers can be directly compared to a reference enhancer in the same animal to detect any functional changes. *Dual-enSERT* also provides a more time-effective and quantitative method for *in vivo* mutagenesis of enhancersthat allows a whole embryo readout of changes in enhancer activity. Another potential application of *dual-enSERT* is genetic labeling of specific cell populations. It is often difficult to isolate a desired cell population with a single genetic driver but using an intersectional strategy with two fluorescent reporters driven by different enhancers can enable labeling and isolation of smaller cellular populations (Daigle et al., 2018; He et al., 2016; Matho et al., 2021).

When testing non-mouse enhancers with *dual-enSERT*, we cannot rule out the potential effects caused by *trans* regulatory divergence between different species, especially if no detectable changes in variant enhancer activity are observed (Kvon et al., 2020). In addition, *Dual-enSERT* has a limited throughput which only allows testing two enhancer alleles per construct. Nevertheless, *dual-enSERT* provides a valuable addition to high-throughput methods such as MPRAs, as changes in enhancer activities are detected *in vivo* in whole live mice and in a native chromosomal context. In summary, our work demonstrates the power of mouse transgenesis by enabling rapid and quantitative comparative *in vivo* testing of disease-linked variants for interpretation of new human genetics findings.

## Acknowledgments

The authors would like to acknowledge the UCI Transgenic Mouse Facility for help in the generation of transgenic mouse lines and Elizabeth Pollina (Washington University at St. Louis) and Daniel Gillam (Harvard) for their generous technical assistance with the nested PCR protocol. This work was made possible, in part, through access to the Genomics Research and Technology Hub Shared Resource of the Cancer Center Support Grant (P30CA-062203) at the University of California, Irvine and NIH shared instrumentation grants 1S10RR025496-01, 1S10OD010794-01, and 1S10OD021718-01. The authors also thank Kvon lab members for their critical reading and suggestions. This work was supported by a National Institutes of Health grant DP2GM149555 (to E.Z.K.). E.W.H. was supported by predoctoral fellowships (F30HD110233, T32NS082174, and T32GM008620) from the National Institutes of Health.

## Author contributions

E.W.H. and E.Z.K. conceived the project. E.W.H., T.A.L., S.H.J., and C.X.C., performed mouse transgenesis experiments. E.W.H. performed single-cell RNA-seq experiments and analyzed the data. J.A.A. performed qPCR experiments. E.W.H. and E.Z.K. wrote the manuscript with input from all authors.

## Data and materials availability

The dual-enSERT vectors described in this study are made available at Addgene. See **Figure S4** for schematics of deposited plasmids. Processed and raw scRNA-seq data are available for download at GEO: GSE244244.

## Materials and Methods

### Ethics statement

This research complies with all relevant ethical regulations. All animal procedures, including those related to the generation of transgenic animals, were conducted in accordance with the guidelines of the National Institutes of Health (NIH) and approved by the Institutional Animal Care and Use Committee at the University of California, Irvine.

### Cloning of *dual-enSERT* constructs

#### *Dual-enSERT-1* plasmid construction

To create *dual-enSERT-1* constructs, we used PCR4-Hsp68::lacZ-H11 plasmid (Kvon et al., 2020) (Addgene #139099) and replaced *lacZ* with *eGFP* or *mCherry* fluorescent reporters using Gibson cloning (Gibson et al., 2009). The resulting constructs contained homology arms targeting the H11 locus and *eGFP* (PCR4-Hsp68::eGFP-H11) or *mCherry* (PCR4-Hsp68::mCherry-H11) fluorescent reporter genes under control of the *hsp68* minimal promoter. Each tested enhancer sequence was cloned into the corresponding *dual-enSERT-1* vector using NotI digestion followed by Gibson assembly methods (NEB, E2611) (Gibson et al., 2009; Osterwalder et al., 2022). Reference allele enhancer sequences for hZRS and hs737 were obtained via PCR cloning from human genomic DNA (Promega, G304A). Primers used for each enhancer sequence are outlined in **Table S2**. All PCR cloning was performed using Q5 High-Fidelity Polymerase (NEB, M0491) or KOD polymerase (Toyobo, #KMM-201). Variant allele sequences for hZRS^404G>A^, hZRS^446T>A^, and hs737^830G>A^ were custom synthesized as gBlocks (Integrated DNA Technologies (IDT)).

#### *Dual-enSERT-2.0* plasmid construction

The bicistronic plasmid was designed as a modified version of the fluorescent *dual-enSERT-1* backbone. First, the cHS4 insulator sequence was cloned from genomic chicken DNA (Zyagen, GC-314) using the primers published in Bhatia et al (Bhatia et al., 2021). To obtain three copies, linker sequences (fragments of the LacZ gene) were added as overhangs to the primers. Enhancer-*hsp68p*-reporter sequences were then PCR amplified from *dual-enSERT-1* plasmids. Gibson-based assembly methods were then used to assemble the finalized plasmid.

#### Dual-enSERT-2.1 and 2.2 plasmid construction

The *dual-enSERT-2.1* and *2.2* plasmids were designed equivalent to *dual-enSERT-2.0*, except with the synthetic insulator replacing three copies of cHS4. To construct the synthetic insulator, each separate insulator sequence was first isolated. The A2 insulator was synthesized as a gBlock using the sequence reported by Liu et al (Liu et al., 2015) while ALOXE3 and cHS4 were cloned from human genomic DNA using primers reported by Raab et al (Raab et al., 2012) and Bhatia et al (Bhatia et al., 2021), respectively. Fusion PCR was performed to obtain the final synthetic insulator fragment from two copies of A2, one copy of ALOXE3, and two copies of cHS4. Enhancer-*hsp68p*-reporter sequences were then PCR amplified from *dual-enSERT-2.0* plasmids. Gibson-based assembly methods were then used to assemble the finalized plasmid.

To streamline the cloning of different enhancers into *dual-enSERT-2.1* plasmids, NotI and AgeI restriction digestion sites were added to the outside of the two enhancer sites via PCR. *Dual-enSERT-2.1* plasmid was digested with NotI (NEB, R3189) and AgeI (NEB, R3552) to create an empty vector without enhancers. To quickly generate different *dual-enSERT-2.1* plasmids, a four-fragment Gibson-based assembly mix was set-up consisting of (i) empty *dual-enSERT-2.1* backbone, (ii) reference enhancer allele, (iii) variant reference allele, and (iv) synthetic insulator. The hZRS and hs737 reference and variant allele sequences tested via *dual-enSERT-2.2* were PCR amplified from *dual-enSERT-1.0* plasmids. To obtain hs932 sequences for *dual-enSERT-2.2*, the hs932^ref^ allele was cloned from human genomic DNA using primers listed in **Table S2** and the hs932^350dupA^ variant allele was synthesized as a gBlock (IDT).

To enable linearisation of the *dual-enSERT-2.1* donor plasmid, PauI (also known as BssHII) sites were added to the outside of the H11 homology arms via PCR. For *dual-enSERT-2.2*, a *dual-enSERT-2.1* plasmid (1 μg) was digested overnight with PauI (NEB, R0199) at 50°C in rCutSmart Buffer. The following day, the reaction was inactivated by incubation at 65°C for 15 min. To end-fill the 3’ overhangs with biotinylated nucleotides, Biotin-11-GTP (100 μM; Jena Bioscience, NU-971-BIOX) and biotin-11-CTP (100 μM; Jena Bioscience, NU-831-BIOX) were added to the reaction with T4 DNA Polymerase (1 unit; NEB, M0203) and incubated at 12°C for 15 min. The reaction was stopped by addition of EDTA (100 mM) and heat inactivated at 75°C for 20 min. Biotinylation of fragments was confirmed by pull-down with streptavidin T1 Dynabeads (Thermo Fisher Scientific, cat. no. 65601).

For all constructs in this study, restriction digestion with SacI and Eco72I (Thermo Fisher, FastDigest), Sanger and/or whole-plasmid (Plasmidsaurus) sequencing were performed to ensure integrity of the vector and enhancer sequences before microinjection. See **Table S3** for complete details of all plasmids created and used in this study.

#### Transgenic mouse generation

All transgenic mice in this study were generated using a CRISPR/Cas9 microinjection protocol, as previously described (Kvon et al., 2016, 2020). Briefly, a mix of (i) Cas9 protein (final concentration of 20 ng/μl; IDT Cat. No. 1074181), (ii) sgRNA (50 ng/μl) and (iii) donor plasmid (7 ng/μl) or linearised fragment (1 ng/μl) in injection buffer (10 mM Tris, pH 7.5; 0.1 mM EDTA) was injected into the pronucleus of FVB embryos. All donor plasmids or fragments were column purified using a PCR purification kit (Qiagen) and eluted into injection buffer before injection. Female mice (CD-1 strain) were used as surrogate mothers. Super-ovulated female FVB mice (7–8 weeks old) were mated to FVB stud males, and fertilized embryos were collected from oviducts. The injected zygotes were cultured in M16 with amino acids at 37°C under 5% CO2 for approximately 2 hr. Afterward, zygotes were transferred into the uterus of pseudopregnant CD-1 females. F0 embryos were either brought to gestation (*dual-enSERT-1*) or collected at E11.5 (*dual-enSERT-2*).

#### Mouse strains and embryo collection

Mice were maintained in standard housing conditions on a reversed 12-hr dark–light cycle with food and water provided *ad libitum*. Time of gestation was identified by the presence of vaginal sperm plugs, indicating E0.5. Pregnant dams were humanely euthanized and embryos carefully removed under brightfield stereoscopes in ice-cold PBS (Cytiva, SH30256.01). Yolk sacs or tail pieces were collected for genotyping. Successful integration events at the H11 locus were determined by PCR using primers described previously (Kvon et al., 2020; Osterwalder et al., 2022).

#### Live fluorescent imaging and quantification

Embryos were imaged in ice-cold PBS in a small petri dish (Greiner Bio-One, #627102) atop a thin layer of 2% gel agarose (Fisher, BP160). Images were taken on a Zeiss V20 stereoscope using a monochromic camera (Axiocam 202, Zeiss), fiber optic light source (Zeiss, CL1500) LED fluorescent laser (X-Cite, Xylis), and fluorescent channels at 488 and 555 nm wavelengths. Single channel images were merged using Zeiss BioLite software. Quantification of fluorescent reporter intensity was performed by importing the original .czi files into Fiji software (Schindelin et al., 2012). Regions of interest were outlined on the variant-driven color channel and then transferred to the reference-driven color channel to measure the mean fluorescence intensities for each region. Because the hsp68 promoter causes leaky reporter activity in the heart (Kvon et al., 2020), the heart was used to control for intrinsic differences between reporter intensity, including as a negative control in two-sided, paired t-tests.

#### Fluorescent-activated cell sorting

After imaging, the anterior portions of *dual-enSERT-1* mouse limb buds carrying two hZRS reporter transgenes were carefully dissected under the fluorescent scope to avoid contamination by the ZPA. Dissected regions from each embryo were pooled separately and then incubated with collagenase II (Gibco, #17101015, 0.2 μL at 100 u/μl) for 10 min at 700 rpm and 37°C with trituration every 5 min with a P200 pipette. Then, 450 μL of 10% FBS (Thermo Fisher, #A3840201) was added and dissociated cells were spun down. Cells were resuspended in 200 μL of 0.04% BSA (Millipore Sigma, #A1595) and filtered using 40 μM P1000 Flowmi cell filters (SP Bel-Art, #136800040). After gating with forebrain tissue as a negative control, mCherry+, eGFP+, and double-positive cell populations were quantified using a FACSAria Fusion Sorter (BD Biosciences). Fisher exact tests were performed between genotypes of double-positive, mCherry+, and eGFP+ cells.

#### Quantitative PCR

Quantitative PCR was performed for *mCherry* and *eGFP* transgenes to determine the copy-number of enhancer-reporter constructs in each *dual-enSERT-1* mouse line (**Fig. S5**). From mouse genomic DNA, *mCherry* or *eGFP* transgenes and an endogenous control of known copy-number were amplified by primers and fluorescent probes (FAM or HEX, IDT), designed using PrimerQuest Tool (IDT) (**Table S2**). Reactions were performed in multiplex and carried out with PrimeTime Gene Expression Master Mix (IDT, 105577) in a C1000 Touch Thermal Cycler with a CFX96 Real-Time System module (Bio Rad, 1845096). Cycle threshold values (Ct) for each amplicon were extracted from .zpcr files using CFX Maestro Software (Bio Rad). Transgene copy numbers (normalized to endogenous control) were calculated using a modified 2−ΔΔCT method in which samples of unknown copy number were compared to positive control samples containing verified single transgene insertions (Livak & Schmittgen, 2001). Only mouse lines with the same copy number of transgenes were compared.

#### Single-cell transcriptomics

A single hindlimb bud from an hZRS^ref^-*mCherry*/hZRS^404G>A^-*eGFP* E11.5 mouse embryo was dissected in ice-cold PBS. A single-cell suspension was obtained using collagenase II, as described above for FACS. Dissociated cells were resuspended in 25 μL of 0.04% BSA before being quantified and inspected for viability using Trypan Blue (Bio-Rad, #1450013). Live cells were counted by hemocytometer (Bio-Rad, #1450011) and loaded at a concentration that would enable recovery of 10,000 nuclei by the Chromium Next GEM Chip G Single Cell Kit 3’ Gene Expression Kit v3.1 (10X Genomics, cat no. 1000127). Captured cells were pair-end sequenced on an Illumina NovaSeq 6000 for approximately 50,000 reads per cell.

To amplify reporter transcripts, nested PCR for mCherry and eGFP transcripts was performed on the prepared cDNA library as described previously, with slight modifications (Pollina et al., 2023). The first PCR included a trimer mix of *mCherry*-and *eGFP*-specific forward primers (mCherry-1; eGFP-1) and an R1-targeting reverse primer (**Table S2**). After bead clean-up (CleanNA, CNGS-0050), a second PCR reaction was performed on the PCR product using a trimer mix targeting *eGFP* and *mCherry* (mCherry-2; eGFP-2) and the same R1 reverse primer (**Table S2**). Resulting DNA was then bead-purified in preparation for the final PCR using the i5:i7 indices from Chromium Next GEM Single Cell 3’ GEM, Library & Gel Bead Kit v3.1 (10X Genomics, cat. no 1000128). All PCR reactions were performed using Q5 polymerase (NEB, M0491). The final indexed and purified DNA was spiked-in for additional sequencing of 5,000 reads per cell.

Fastq files were aligned to a modified mm39 genome assembly (Ensembl, GCA_000001635.9) that included *mCherry* and *eGFP* sequences and barcodes were counted to generate a count matrix using CellRanger (10x Genomics, Cell Ranger 3.1.0). Count matrix data were analyzed using the Seurat R package, version 4 (Hao et al., 2021). A SeuratObject was created and quality control performed to exclude cells with greater than 5% mitochondrial DNA and with UMI counts between 2500 and 8000 per cell. Transcriptome data were normalized, scaled, and variably-expressed genes (n=2000) were utilized for principal component analysis. An elbow plot was produced to calculate the number of dimensions. Because the developing E11.5 limb bud is highly proliferative, cell cycle genes (S and G2M genes from Seurat) were regressed out to enhance detection of cell clusters based on spatial gene expression. Nearest neighbor, unsupervised clustering and UMAP analysis were then performed (dimensions = 13, resolution = 0.5). Cell types were identified from the resulting clusters using well-defined marker genes (Desanlis et al., 2020; Yokoyama et al., 2017). FeaturePlot was utilized to produce the overlaid plots of *mCherry* and *eGFP* expression. The DotPlot function was used to generate expression data by each cell cluster. To determine the topmost upregulated genes within the normal and ectopic ZRS domains, cells were selected for expression of the reporter transcripts: either *mCherry* > 2 (normal ZRS) or *eGFP* > 2 (variant ZRS), or inactive (remaining cells). Differential gene expression was defined by comparing cell populations using the FindMarkers function and the heatmap was downsampled (n=100) for easier visualization. All barcode meta-data, including cell annotations, are reported in **Table S4**.

#### Statistics and reproducibility

Statistical analyses were performed using R version 4.3.1 and Microsoft Excel version 16.79.1. Experimental parameters, including the number of embryos, statistical tests, and significance are reported in the text, figures and/or figure legends. *P* values less than 0.05 were considered significant. All bar graphs are shown as mean ± SEM. Raw sequencing data were analyzed on the UCI high-performance computing cluster.

## Supplementary Figures

**Supplementary Figure 1:**
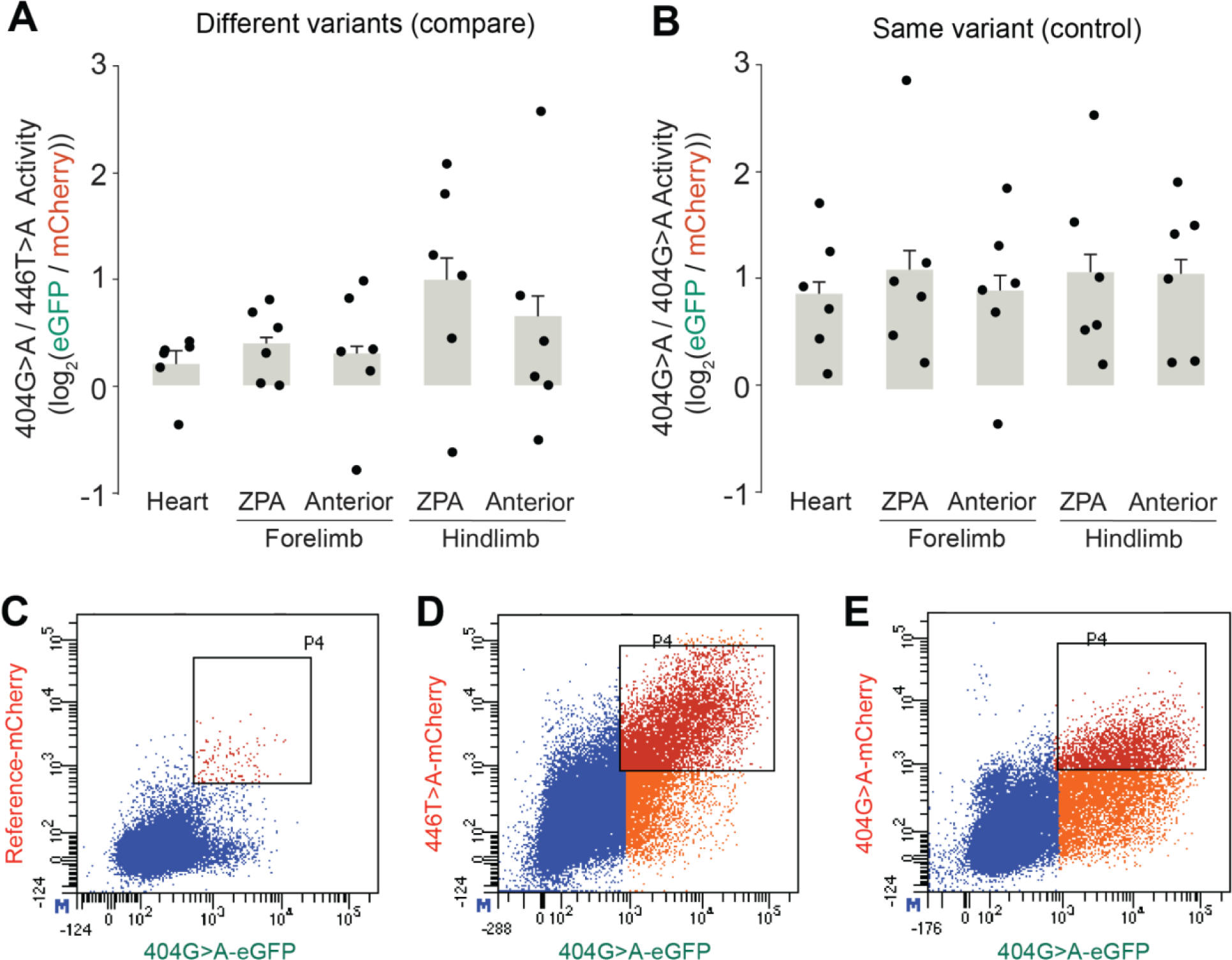
Quantitative analysis of ZRS variant allele-driven reporter intensities. (**A**) Quantitative plot for fold-change (log2) difference in fluorescent reporter intensity between 404G>A and 446T>A variant alleles from hZRS^446T>A^-*mCherry*/hZRS^404G>A^-*eGFP* embryos at E11.5. Two-sided paired t-tests: Forelimb ZPA, *P =* ns; Forelimb Anterior, *P =* ns; Hindlimb ZPA, *P =* ns; Hindlimb Anterior, *P =* 0.0387. (**B**) Quantitative plot for fold-change (log2) difference in fluorescent reporter intensity between the same 404G>A variant allele from hZRS^404G>A^-*mCherry*/hZRS^404G>A^-*eGFP* embryos at E11.5. Two-sided paired t-tests: Forelimb ZPA, *P =* ns; Forelimb Anterior, *P =* ns; Hindlimb ZPA, *P =* ns; Hindlimb Anterior, *P =* ns. Data represented as mean ± SEM for all plots. (**C**) Representative flow cytometry plots for anterior portion of E11.5 hindlimbs from *hZRS^ref^-mCherry/hZRS^404G>A^-eGFP.* (**D**) Sample flow cytometry plots for anterior portion of E11.5 hindlimbs from *hZRS^446T>A^-mCherry/hZRS^404G>A^-eGFP.* (**E**) Example flow cytometry plots for anterior portion of E11.5 hindlimbs from *hZRS^404G>A^-mCherry/hZRS^404G>A^-eGFP.*

**Supplementary Figure 2:**
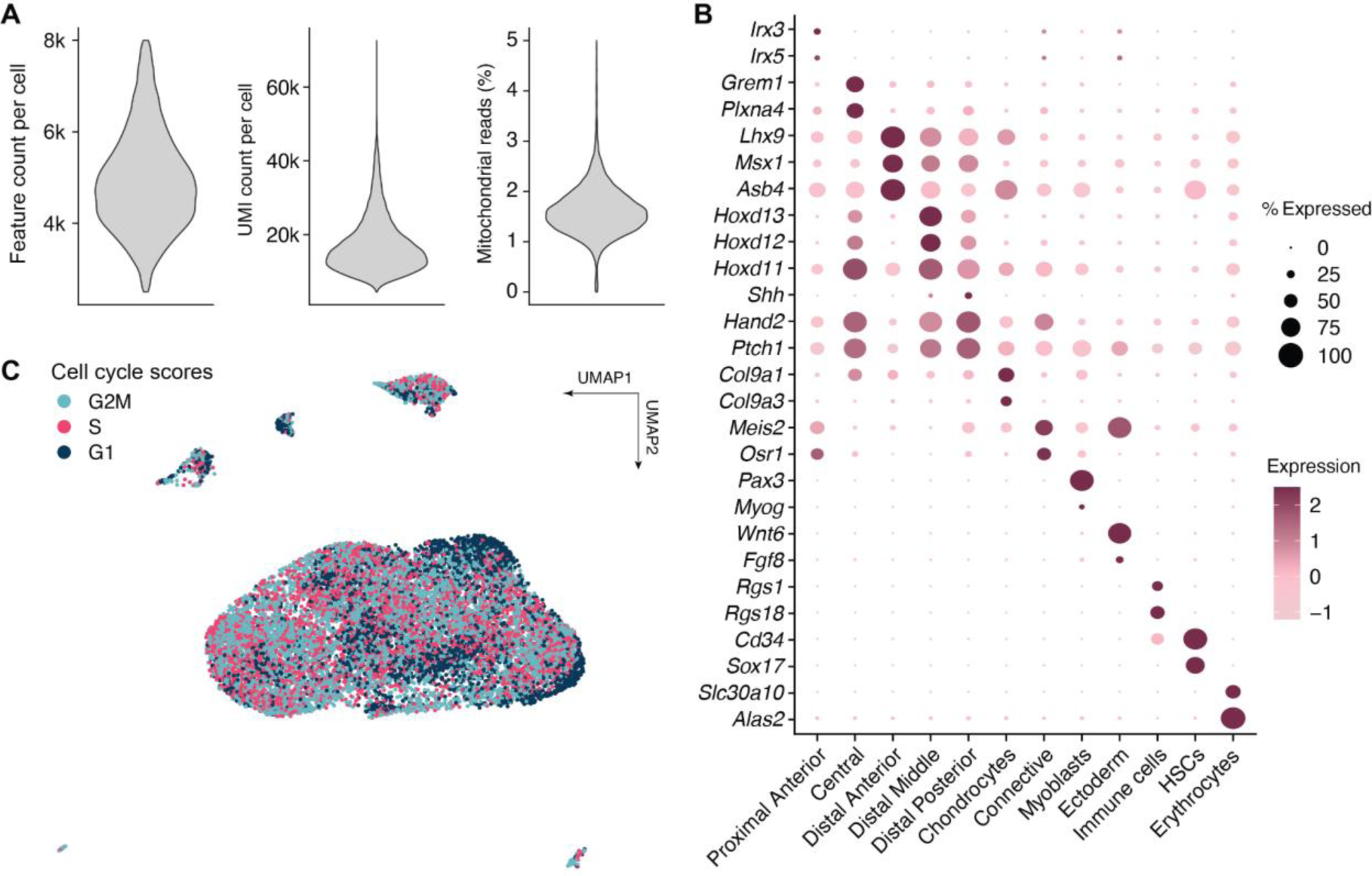
Quality control for scRNA-seq from E11.5 hindlimb of an hZRS^ref^-*mCherry*/hZRS^404G>A^-*eGFP* embryo. **(A)** Plots depicting feature and UMI counts and percent mitochondrial genes expressed. **(B)** DotPlot of cell type marker expression across all clusters. **(C)** UMAP plot depicting cell cycle scores assigned by regression analysis.

**Supplementary Figure 3:**
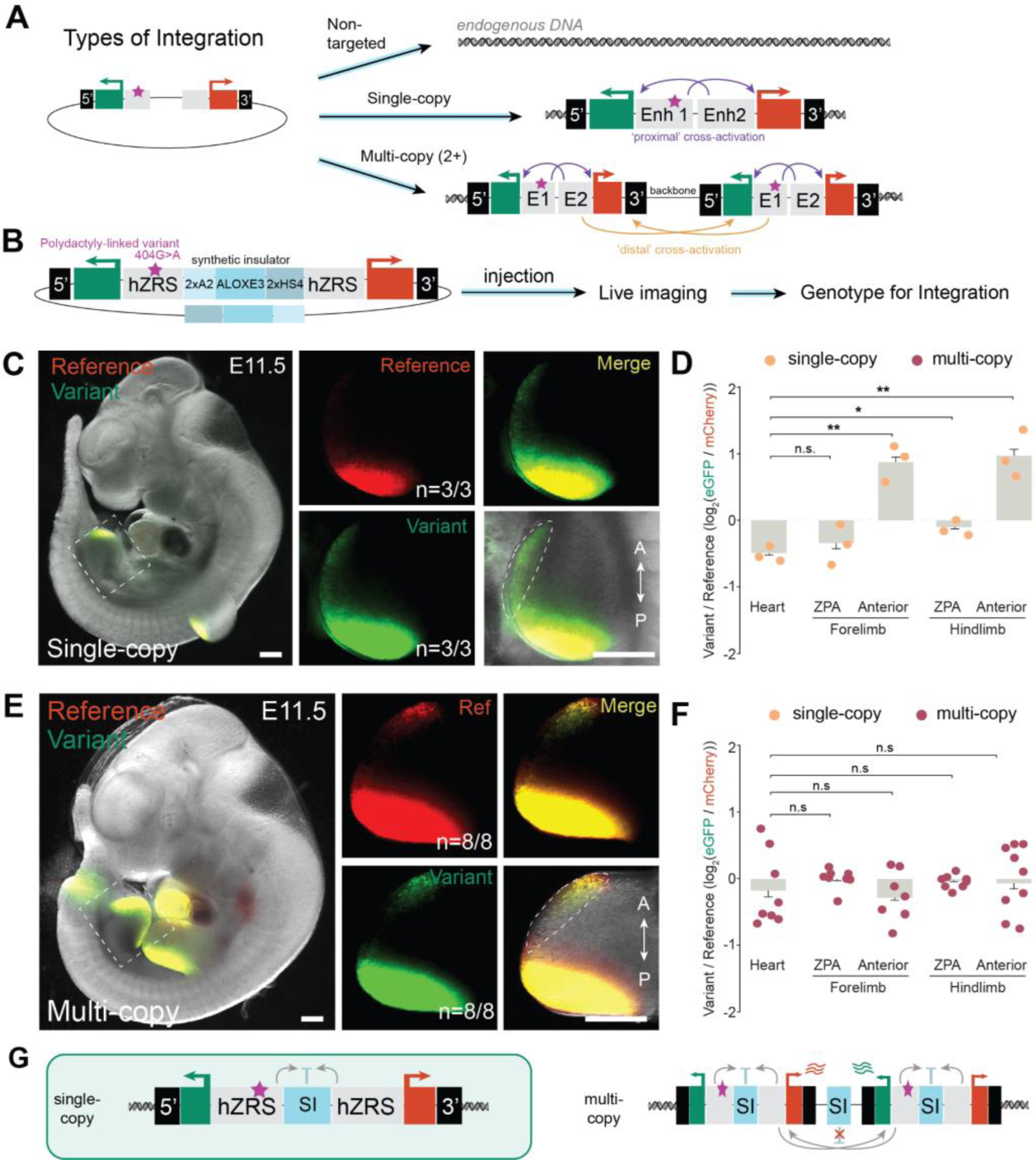
Optimization of a strong, synthetic insulator for *dual-enSERT-2.1* with single and multi-copy integrations. **(A)** Types of integrations of donor plasmid into the H11 locus and the potential scenarios of enhancer cross-activation. (**B**) Experimental approach for injection of ZRS^ref^-*mCherry*/SI/ZRS^404G>A^-*eGFP* plasmid into mouse embryos with collection and genotyping. (**C**) Sample images of single-copy embryos injected ZRS^ref^-*mCherry*/SI/ZRS^404G>A^-*eGFP* construct with hindlimb highlighted by dashed box and higher-resolution images on right. (**D**) Plots quantifying fold-change (log2) difference in reporter intensity for single-copy integrants. Two-sided paired t-test: Single-copy: Forelimb ZPA, *P =* ns; Forelimb Anterior, *P =* 0.00126; Hindlimb ZPA, *P =* 0.0110; Hindlimb Anterior, *P =* 0.00235. Data represented as mean. (**E**) Sample images of multi-copy embryos injected ZRS^ref^-*mCherry*/SI/ZRS^404G>A^-*eGFP* construct with hindlimb highlighted by dashed box and higher-resolution images on right. (**F**) Plots quantifying fold-change (log2) difference in reporter intensity for multi-copy integrants. Two-sided paired t-tests. Multi-copy: Forelimb ZPA, *P =* ns; Forelimb Anterior, *P =* ns; Hindlimb ZPA, *P =* ns; Hindlimb Anterior, *P =* ns. Data represented as mean ± SEM. Points representing single-(n=3) and multi-copy (n=8) integrants are in orange and maroon, respectively. (**G**) Schematic summary of results for *dual-enSERT-2.1* with the synthetic insulator only preventing cross-activation when integrated as a single-copy.

**Supplementary Figure 4.**
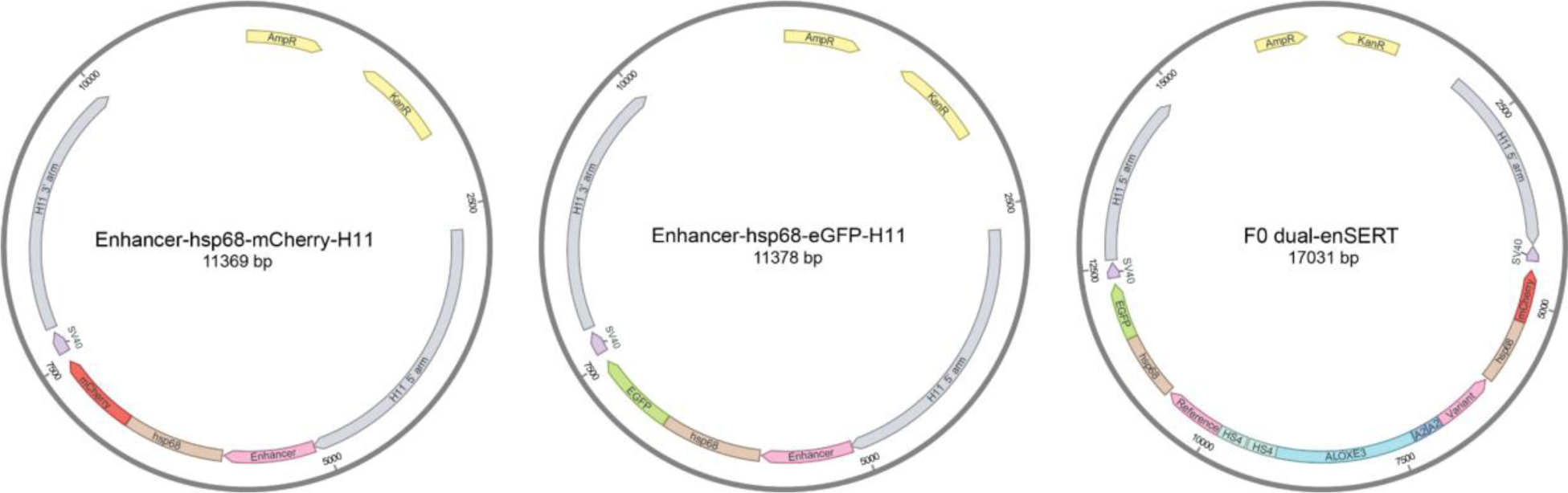
Schematic representation of plasmids deposited by this study. Note that deposited versions of these plasmids are empty backbones without any enhancer sequences.

**Supplementary Figure 5:**
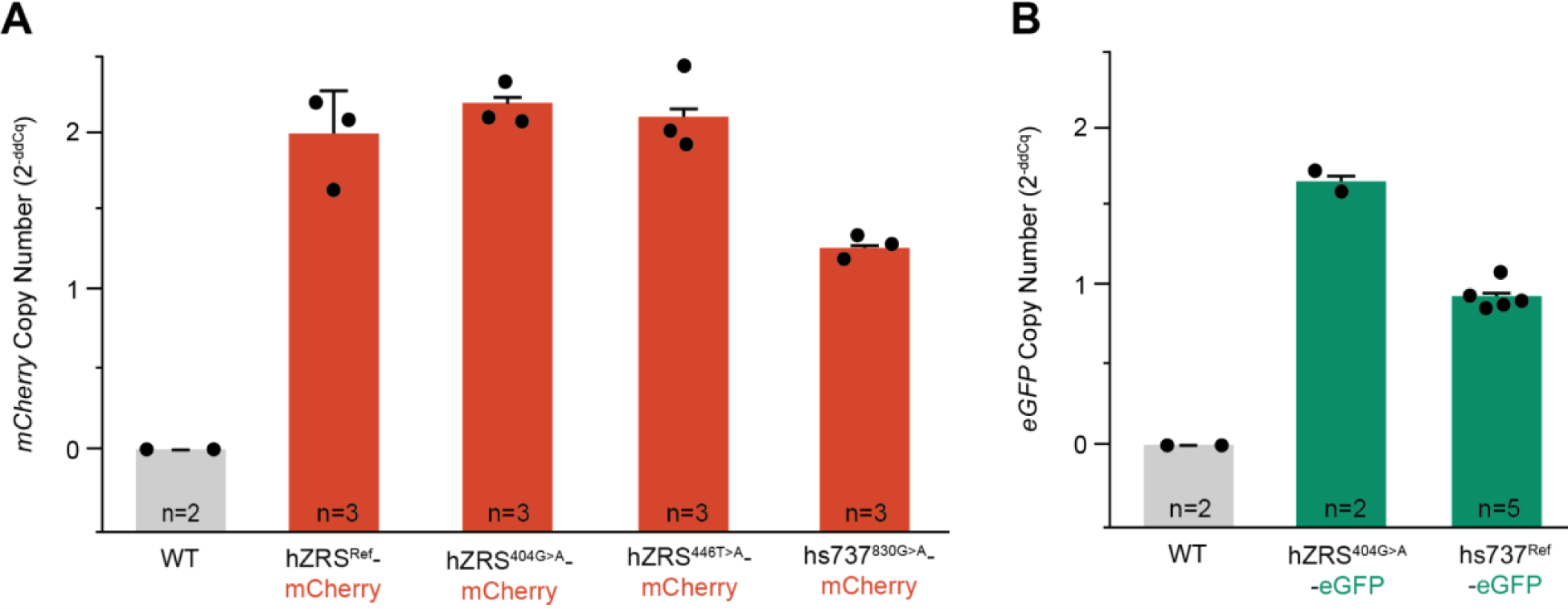
Quantification of the number of enhancer-reporter constructs integrated into each mouse line by qPCR. **(A, B)** Quantitative PCR plot for mCherry (A) and eGFP (B) copy number across *dual-enSERT-1* lines. Note equivalent copy number for each enhancer allele (i.e., two for hZRS and one for hs737).

## Supplementary Tables

Note: Supplementary Table S4 is provided in a separate Excel spreadsheet due to its size.

**Table S1.** Genotyping and cross-activation results for *dual-enSERT-2* constructs.

| Construct | Genotype | Cross-activation | Insulation | Total |
| --- | --- | --- | --- | --- |
| hZRS<sup>ref</sup>-<br>*mCherry*/3xHS4/hZRS<sup>404G>A</sup>-<br>*eGFP*<br>(*dual-enSERT-2.0*) | Single-copy | 5/5 | 0/5 | 5 |
|  | Multi-copy | 11/11 | 0/11 | 11 |
|  | Total F0 at H11 | 16/16 | 0/16 | 16 |
| hZRS<sup>ref</sup>-*mCherry*/SI/<br>hZRS<sup>404G>A</sup>-*eGFP*<br>(*dual-enSERT-2.1*) | Single-copy | 0/3 | 3/3 | 3 |
|  | Multi-copy | 8/8 | 0/8 | 8 |
|  | Total F0 at H11 | 8/11 | 3/11 | 11 |
| B-hZRS<sup>ref</sup>-*mCherry*/SI/<br>hZRS<sup>404G>A</sup>-*eGFP*-B<br>(*dual-enSERT-2.2*) | Single-copy | 0/5 | 5/5 | 5 |
|  | Multi-copy | 1/1 | 0/1 | 1 |
|  | Total F0 at H11 | 1/6 | 5/6 | 6 |
| B-hs737<sup>ref</sup>-*eGFP*/SI/<br>hZRS<sup>830G>A</sup>-*mCherry*-B<br>(*dual-enSERT-2.2*) | Single-copy | 0/3 | 3/3 | 3 |
|  | Multi-copy | 0/0 | 0/0 | 0 |
|  | Total F0 at H11 | 0/3 | 3/3 | 3 |
| B-hs932<sup>ref</sup>-*eGFP*/SI/<br>hs932<sup>350dupA</sup>-*mCherry*-B<br>(*dual-enSERT-2.2*) | Single-copy | 0/4 | 4/4 | 4 |
|  | Multi-copy | 0/0 | 0/0 | 0 |
|  | Total F0 at H11 | 0/4 | 0/4 | 4 |

**Table S2.**
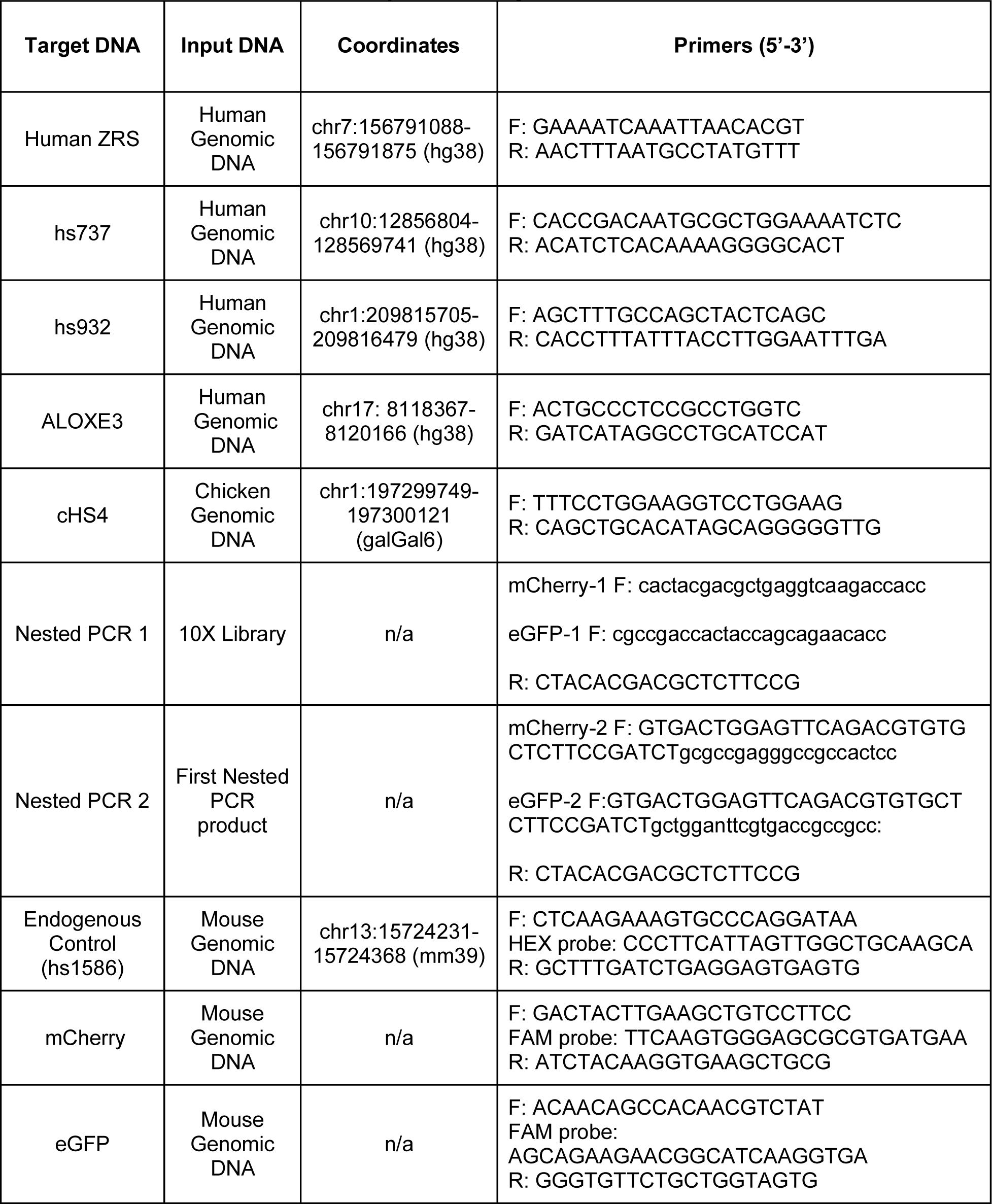
Primers used in this study for cloning, nested PCR, and qPCR.

**Table S3.** Plasmids created and deposited by this study.

| Plasmid name | *dual-enSERT* version |
| --- | --- |
| PCR4-Hsp68::mCherry-H11 (empty vector) | *dual-enSERT-1.0* |
| PCR4-hZRS<sup>ref</sup>-Hsp68::mCherry-H11 | *dual-enSERT-1.0* |
| PCR4-hZRS<sup>404G>A</sup>-Hsp68::mCherry-H11 | *dual-enSERT-1.0* |
| PCR4-hZRS<sup>446T>A</sup>-Hsp68::mCherry-H11 | *dual-enSERT-1.0* |
| PCR4-hs737<sup>830G>A</sup>-Hsp68::mCherry-H11 | *dual-enSERT-1.0* |
| PCR4-Hsp68::eGFP-H11 (empty vector) | *dual-enSERT-1.0* |
| PCR4-hZRS<sup>404G>A</sup>-Hsp68::eGFP-H11 | *dual-enSERT-1.0* |
| PCR4-hs737<sup>ref</sup>-Hsp68::eGFP-H11 | *dual-enSERT-1.0* |
| PCR4-Hsp68::mCherry-3xHS4-Hsp68::eGFP-H11(empty vector) | *dual-enSERT-2.0* |
| PCR4-hZRS<sup>ref</sup>-Hsp68::mCherry-3xHS4-hZRS<sup>404G>A</sup>-Hsp68::eGFP-3xHS4-H11 | *dual-enSERT-2.0* |
| PCR4-Hsp68::mCherry-SI-Hsp68::eGFP-H11-SI (empty backbone) | *dual-enSERT-2.1* |
| PCR4-hZRS<sup>ref</sup>-Hsp68::mCherry-SI-hZRS<sup>404G>A</sup>-Hsp68::eGFP-SI-H11 | *dual-enSERT-2.1* |
| PCR4-Hsp68::mCherry-SI-Hsp68::eGFP-H11 (empty vector) | *dual-enSERT-2.2* |
| PCR4-hZRS<sup>ref</sup>-Hsp68::mCherry-SI-hZRS<sup>404G>A</sup>-Hsp68::eGFP-H11 | *dual-enSERT-2.2* |
| PCR4-hs932<sup>350dupA</sup>-Hsp68::mCherry-SI-hs932<sup>ref</sup>-Hsp68::eGFP-H11 | *dual-enSERT-2.2* |

**Table S4. [see separate Excel file]**

**Barcode meta-data for single-cell transcriptomics of dual-enSERT E11.5 hindlimb with human reference and variant ZRS alleles**

